# Cognitive Baseline and Stimulation Site Shape the Effects of tDCS on Verbal Fluency in Older and Younger Adults

**DOI:** 10.1101/2025.08.22.671683

**Authors:** A. Yucel, A. K Martin

## Abstract

Verbal fluency is a core measure of language and executive function often used in cognitive research. Transcranial direct current stimulation (tDCS) has been investigated for its potential to enhance verbal fluency; however, the results are variable. Discrepancies may indicate variations in electrode montage, stimulation site, measured fluency type, and individual cognitive profiles. In this preregistered, double-blind, sham-controlled study, 72 younger and 72 older adults received anodal tDCS to either the left inferior frontal gyrus (left IFG) or the left temporoparietal junction (left TPJ) while completing phonemic and semantic fluency tasks. Anodal stimulation of the left TPJ significantly improved phonemic fluency compared to sham (*F*(1,71) = 4.49, p = .038), with no effects observed for semantic fluency or left IFG stimulation. Additionally, stimulation response in the TPJ group was predicted by fluid intelligence, with lower scores associated with greater benefit (F(1,70) = 7.80, p = .007). Finally, stimulation to the left IFG improved response initiation but impaired sustained energization over the entire minute. These results suggest that the efficacy of focal tDCS for enhancing verbal fluency depends on both stimulation site and task demands, with the left TPJ showing selective benefits for phonemic fluency. Importantly, individual differences in fluid intelligence, but not age, moderated stimulation response, highlighting the role of cognitive capacity in neuromodulation outcomes. Together, these findings demonstrate a nuanced relationship between targeted excitation of the language network and verbal fluency, emphasizing the need for individualized approaches in cognitive enhancement interventions.

## Introduction

Verbal fluency, defined as the capacity to rapidly generate words based on designated phonemic or semantic criteria, serves as a fundamental aspect of language processing and is frequently employed as a measure in cognitive and clinical evaluations (Spreen & Strauss, 1998; Shao et al., 2014) due to its sensitivity to decline in healthy and pathological ageing. Impairments in lexical access and executive control are important factors affecting verbal fluency, leading to reduced communicative effectiveness (James & Burke, 2000; Henry & Crawford, 2004). This has resulted in an increased scientific interest in identifying interventions that might support verbal fluency across the healthy lifespan or slow its decline. One such approach is transcranial direct current stimulation (tDCS), a non-invasive neuromodulatory technique that alters cortical excitability by applying weak, low-intensity electrical currents to the scalp. Evidence indicates that tDCS may improve language performance (Monti et al., 2013); however, it remains uncertain whether focal stimulation reliably improves verbal fluency in healthy adults with studies limited by a focus on one brain region or age group (Price et al., 2015; Westwood et al., 2017). Moreover, investigation into the role of factors such as baseline function, timing of effects, and direct comparisons of phonemic and semantic fluency is lacking. Thus, this study investigates the effect of anodal tDCS applied to two key hubs of the left-lateralized language network on phonemic and semantic fluency in young and older adults. Moreover, we investigate the role of baseline fluid intelligence and energization demands on stimulation response.

Language function, particularly verbal fluency, has become a growing interest in tDCS research, especially in older adults, due to its reliance on a distributed fronto-temporo-parietal network and its susceptibility to age-related cognitive decline (Fertonani & Miniussi, 2017; Bikson et al., 2016). Several studies have reported facilitation of picture naming and word retrieval following anodal stimulation to the left inferior frontal gyrus (left IFG) in both healthy young adults and clinical cohorts (Iyer et al., 2005; Sparing et al., 2008; Fertonani et al., 2010; Baker et al., 2010; Perceval et al., 2017; Yucel et al, 2025). While these findings suggest some promise for tDCS as a language-enhancing tool, the majority of investigations have focused exclusively on frontal cortical targets, often overlooking posterior language-related regions (Westwood & Romani, 2017). One such region is the left temporoparietal junction (left TPJ), a key hub implicated in lexical integration, semantic retrieval, and automatic access to stored verbal representations - mechanisms engaged in verbal fluency tasks, requiring people to rapidly and efficiently produce words based on defined criteria. (Wang & Bastiaansen, 2012; Hirshorn & Thompson-Schill, 2006; Scott et al., 2000). A more comprehensive understanding of how stimulation to these different regions affects language performance is necessary to identify specific functional contributions of anterior and posterior language regions. Such knowledge could then be used to design and apply targeted neuromodulatory interventions, particularly in ageing populations.

A notable strength of the present study is the use of focal tDCS, which constrains current flow to the intended cortical site and thereby achieves greater spatial specificity compared to the more traditional sponge-electrode montages that produce diffuse current spread (Datta et al., 2010; Stagg & Nitsche, 2011; Jacobson et al., 2012; Niemann et al., 2024). This methodological refinement is particularly important in the study of verbal fluency, where prior investigations have largely employed conventional montages and thus limited the precision of causal inferences regarding the contribution of specific brain regions (e.g., Iyer et al., 2005; Cattaneo et al., 2011; Westwood et al., 2017). Verbal fluency itself is not a unitary construct but can be divided into phonemic fluency—requiring word production based on initial letters and strongly linked to executive processes supported by frontal regions—and semantic fluency, which depends on retrieval within a semantic category and is more closely associated with temporoparietal regions implicated in semantic memory (Troyer et al., 1997; Birn et al., 2010).

Neuroimaging and lesion evidence converge on the view that both anterior and posterior language regions play critical roles in supporting these fluency processes (Binder et al., 2009; Vigneau et al., 2006). Yet, the majority of tDCS studies have targeted only a single site, most often the left inferior frontal gyrus (IFG) or adjacent prefrontal areas (Pisoni et al., 2018; Ehlis et al., 2016; Penolazzi et al., 2013; Meinzer et al., 2012; Vannorsdall et al., 2012; Cattaneo et al., 2011). By adopting a focal stimulation montage that separately targets both the left IFG and the left temporoparietal junction (TPJ), the present study moves beyond these limitations, providing a more precise test of the relative causal contributions of anterior versus posterior language regions to phonemic and semantic fluency.

Another critical issue in the existing literature is the limited understanding of how age influences the effects of tDCS on language-related performance. While most of the existing evidence comes from studies involving young adults (Pisoni et al., 2018; Penolazzi et al., 2013; Cattaneo et al., 2011; Cerruti & Schlaug, 2009), it remains unclear whether these effects consistent to older populations who differ in terms of current cognitive capacity and cortical structure. However, ageing is associated with structural and neurochemical changes that may reduce the brain’s responsiveness to stimulation, including cortical thinning, disrupted functional connectivity, and these changes are thought to reduce the effectiveness of tDCS by limiting neuroplastic potential, particularly in prefrontal regions involved in executive aspects of language processing. (Fujiyama et al., 2016; Li et al., 2015; Miniussi et al., 2013). However, other research suggests that tDCS may be particularly effective in older adults, potentially because stimulation can compensate for age-related decline and thus provide greater scope for improvement (Berryhill & Jones, 2012). By including both younger and older participants, the present study is uniquely positioned to examine how age influences the efficacy of focal tDCS in modulating verbal fluency.

Temporal aspects of task performance represent another dimension through which stimulation effects may be observed. Verbal fluency tasks typically show a characteristic temporal pattern, with most responses produced during the first 15 seconds of the trial, followed by a marked reduction in retrieval rate across the remaining 45 seconds (Möckel et al., 2015; Venegas & Mansur, 2011; Hirshorn & Thompson-Schill, 2006). This decline over time has been linked to the cognitive process of energization, which involves the initiation and sustained effort required to maintain performance throughout the task (Robinson et al., 2012). This initial burst is presumed to reflect more automatic lexical access, while later responses depend increasingly on effortful, controlled search processes. Deficits in this early-phase output have been observed in populations with frontal dysfunction (Stuss & Alexander, 2007; Barker et al., 2018), suggesting that frontal regions such as the left IFG play a disproportionate role in the initiation of verbal responses. Given that tDCS modulates cortical excitability, we hypothesized that stimulation over the left IFG would selectively increase word production in the early segment of the task, whereas stimulation over the left TPJ, which is associated with semantic retrieval, would have more sustained effects over time.

Therefore, the current preregistered study (https://osf.io/67g859) aims to provide a comprehensive investigation into the effects of focal tDCS on phonemic and semantic verbal fluency, by targeting two key hubs of the left-lateralised language network, the left IFG and left TPJ. Importantly, stimulation effects are assessed in both healthy younger and older adults, allowing for analysis of age-related differences in neuromodulatory response. Additionally, by separately analyzing early and late phases of fluency output, the study examines whether stimulation interacts with different cognitive stages of lexical retrieval, namely automatic retrieval versus controlled search. This design enables a comprehensive examination of how focal tDCS affects language production as a function of stimulation site, cognitive process, and age, and provides implications for targeted interventions in healthy ageing and clinical language rehabilitation.

## Methods

The study design and all analyses were preregistered online (https://osf.io/67g859)

### Participants

A power analysis was conducted via G*Power software (Faul et al., 2007). To obtain a medium effect size (η² = 0.06, f = 0.25) with an alpha level of 0.05 and a power of 0.80, 36 participants per condition were needed, resulting in a total sample size of 144 participants (36 younger left IFG, 36 younger left TPJ, 36 older left IFG, 36 older left TPJ). Participants were neurologically healthy individuals who were native English speakers and had no prior experience with tDCS. The study included two age groups: older adults aged 55 to 85 (n = 72) and younger adults aged 18 to 30 (n = 72). Young and older adults were stratified to receive stimulation to either the left IFG or the left TPJ in a sham-controlled, repeated-measures design. The young left IFG group (25 females, 11 males), young left TPJ group (28 females, 8 males), older left IFG group (21 females, 15 males), and older left TPJ group (27 females, 9 males) showed comparable gender distributions (See Table 1 for the detailed demographic information of the participants). The older adults were selected from the University of the Third Age (U3A) and the University of Kent Research Hub groups. Upon completing the study, the older adults received a £15 compensation for their time and efforts. The younger adults were University of Kent students and from the surrounding community and were awarded course credits or £15.

**Table 1.**
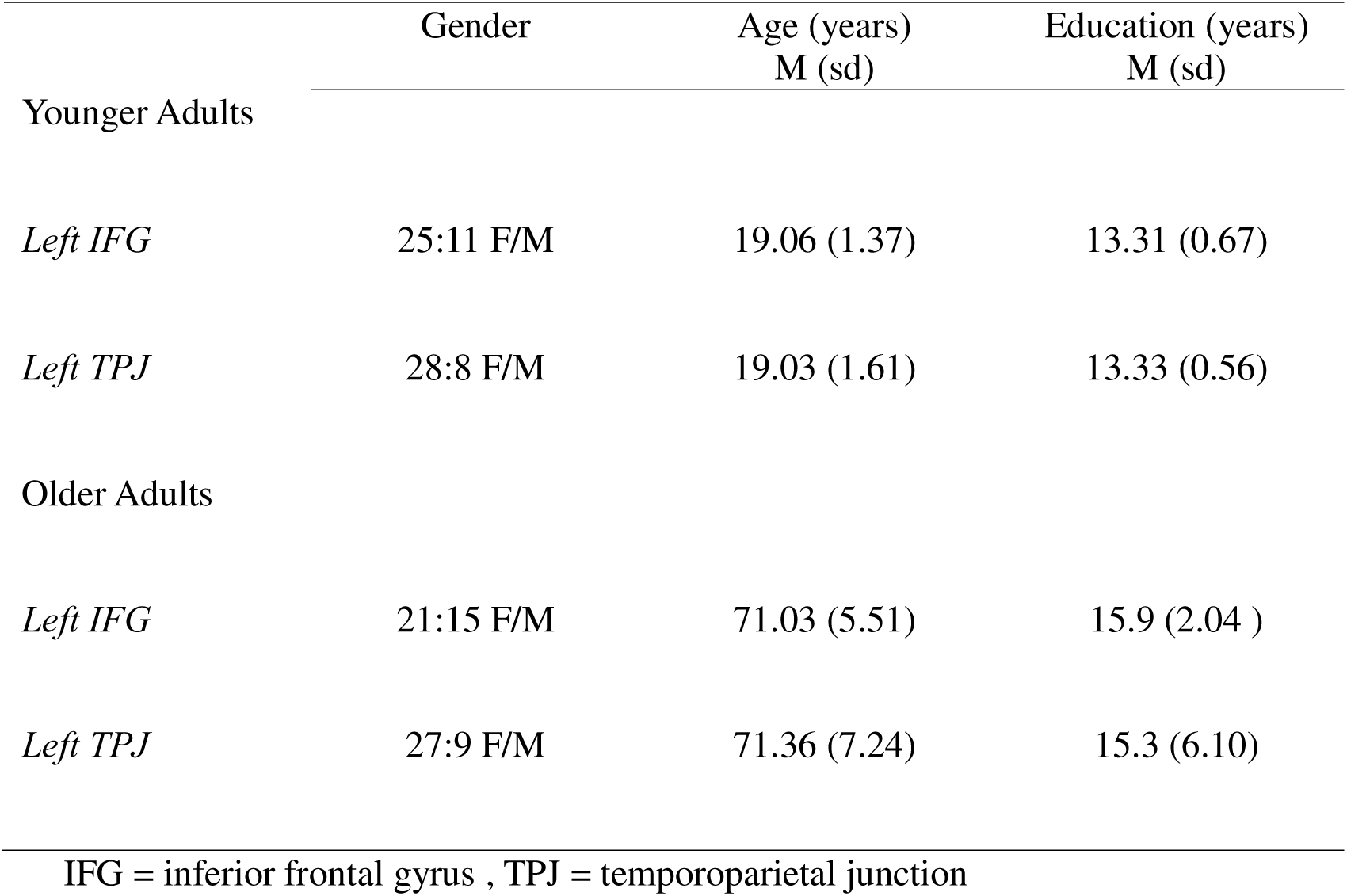
Demographic details for younger and older adults at both the left IFG and the left TPJ stimulation sites.

|  | Gender | Age (years)<br>M (sd) | Education (years)<br>M (sd) |
| --- | --- | --- | --- |
| Younger Adults |  |  |  |
| *Left IFG* | 25:11 F/M | 19.06 (1.37) | 13.31 (0.67) |
| *Left TPJ* | 28:8 F/M | 19.03 (1.61) | 13.33 (0.56) |
| Older Adults |  |  |  |
| *Left IFG* | 21:15 F/M | 71.03 (5.51) | 15.9 (2.04 ) |
| *Left TPJ* | 27:9 F/M | 71.36 (7.24) | 15.3 (6.10) |

IFG = inferior frontal gyrus , TPJ = temporoparietal junction

The study strictly adhered to standard tDCS safety criteria for exclusions, such as a history of seizures, strokes and the presence of metallic items in the head. Participants were neurologically healthy without current or historical mental health conditions. None of the participants disclosed the use of medications that are known to interfere with the effects of tDCS, such as antidepressants and anxiolytics. Each participant provided written informed consent, and the project received official clearance from the Human Research Ethics Committee [ID: 202216587800677857] from The School of Psychology at the University of Kent.

### Verbal Fluency Task

Participants completed two verbal fluency tasks during each stimulation session: semantic and phonemic fluency. In the semantic fluency task, participants were instructed to generate as many words as possible within one minute that belonged to a specified category. For example, in the “animals” category, valid responses might include “dog” and “cat.” Each session included two semantic categories (e.g., “animals,” “fruits and vegetables,” “musical instruments,” or “vehicles”), selected from established norms to ensure general familiarity and balance across sessions (Adams et al., 1989; Acevedo et al., 2000; Kavé, 2005). Categories were counterbalanced across stimulation conditions to control for content effects. In the phonemic fluency task, participants were instructed to generate as many words as possible within one minute that began with a specified initial letter. For example, for the letter “C,” acceptable responses might include “car” and “cold.” Across sessions, participants completed two letter-based fluency tasks. The letter pairs *M–T* (word-starting frequencies: M = 2048, T = 2031) and *G–L* (G = 1220, L = 1206) were selected from Wolfram’s (2015) lexical frequency norms to create comparable task versions differing in difficulty (M–T = easier; G–L = harder). Each participant completed one easy and one hard letter-pair task across the two sessions, with the order of pair assignment counterbalanced across sham and anodal stimulation conditions. Repetitions of the same word root (e.g., “go,” “going,” “gone”) were not counted as separate responses, and proper nouns (e.g., “Germany” “Gareth”) were excluded from scoring. Phonemic fluency tasks always followed the semantic fluency tasks in each session. For statistical analyses, scores from the two versions were combined to provide a single measure of verbal fluency performance per session. We also measured performance in the first 15 and last 45 second time windows, to assess response initiation and energization (as per Barker *et al*, 2018).

### Matrix reasoning item bank (MaRs-IB) task

The Matrix Reasoning Item Bank (MaRs-IB; Chierchia et al., 2019) was used to establish fluid intelligence, to evaluate its predictive value for stimulation effects on verbal fluency. Fluid intelligence is thought to at least partially underpin many executive processes (Duncan et al., 2017), including those relevant to verbal fluency in healthy older adults (Shao et al., 2014). Three by three matrices were used to represent each MaRs-IB pattern. Along with eight other cells with abstract shapes, there is one empty cell on the lower right-hand side of the matrix. The four choice arrays displayed below the matrix’s line required participants to complete the pattern by filling in the missing shape. In order to determine what the missing shape was in relation to the other shapes in the matrix, participants had to decipher the pattern. The MaRs-IB task consisted of 50 patterns and a total score was used in the analyses.

### Focal transcranial direct current stimulation

A battery-powered, one-channel direct current stimulator (DC-Stimulator Plus, NeuroConn, Ilmenau, Germany) was used to deliver the tDCS. The focal tDCS electrode montage consisted of two conductive rubber electrodes: a small disc-shaped centre electrode with a diameter of 2.5 cm and a ring-shaped return electrode with an inner diameter of 7.5 cm and an outer diameter of 10 cm. The temporoparietal junction (CP5 spot of the 10–20 EEG system) or the left inferior frontal gyrus (FC5 spot of the 10–20 EEG system) was the position of the centre electrode. In both stimulation settings, the current was gradually increased to 1 mA over a period of 5 seconds. After a 20-minute anodal transcranial direct current stimulation (tDCS) session or a 40-second sham tDCS session, it remained steady until decreasing by over 5 seconds. Producing a tactile feeling comparable to active stimulation and aiding in bliniding participants to the stimulation received. Using the DC stimulator’s “study mode,” we used a predetermined code that activates either active or sham tDCS, ensuring investigator blinding. The codes were allocated by a researcher who was not involved in the experiment. Figure 2 presents the current modelling for both the left inferior frontal gyrus and the right temporoparietal junction. The simulation of current flow was predicated on a reliable 1 mm MNI152 T1 standard brain of a young adult. The current modelling was conducted using SimNIBS version 4.1 (Thielscher et al., 2015). The normal component of the electric field, which is orthogonal to the cortical surface, is deemed the most physiologically relevant for impacting neuronal activity (Radman et al., 2007).

**Figure 1.**
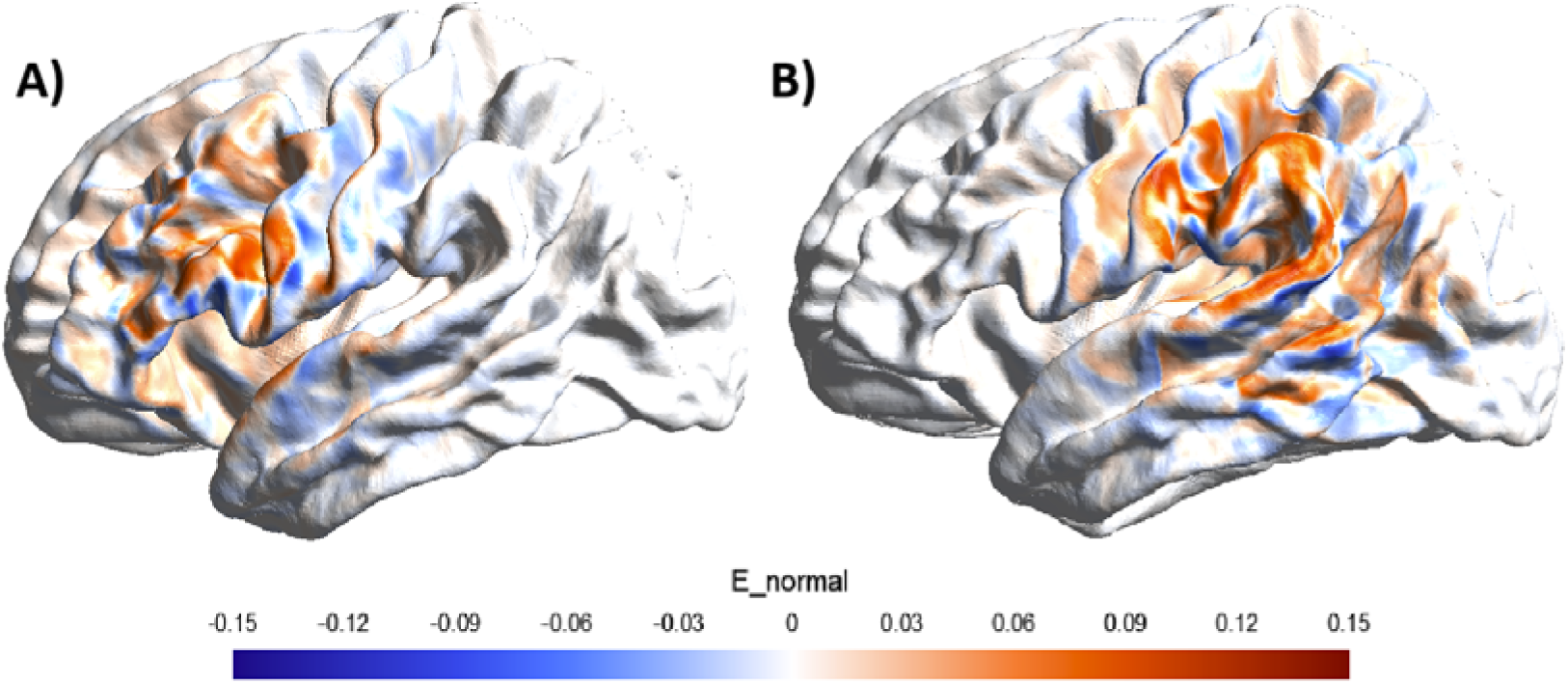
Surface rendering of the distribution and intensity of the estimated electric field for the two target sites with anodal stimulation. A) Left inferior frontal gyrus, B) Left temporoparietal junction. We present the normal component (adapted from Yucel, 2025).

### Mood, adverse effects, and blinding

The mood was assessed using the Visual Analogue Mood Scale (VAMS) before and after each session (Folstein & Luria, 1973). The VAMS measured present positive and negative emotional states using visual analogue scales from 0 to 100, encompassing the following states: afraid, confused, sad, angry, energetic, tired, happy, and tense. Higher scores correspond to more intensity. For each mood, the post-stimulation scores are subtracted from the pre-stimulation scores. Mood is scored into two categories: positive mood (characterised by energy and happiness) and negative mood (fear, confusion, sadness, anger, fatigue, and tension). The deducted scores for each mood and the total scores for positive and negative moods have been obtained for analysis.

Adverse effects were evaluated through the use of a self-report questionnaire created by Brunoni et al. (2011). The participants assessed the intensity (1 = absent, 2 = mild, 3 = moderate, 4 = severe) and occurrence of several possible adverse effects, including headache, neck pain, scalp pain, tingling, itching, burning sensation, skin redness, sleepiness, trouble with concentration, and sudden mood changes. A total adverse effects score was calculated and used in the analysis. Participant blinding was tested at the end of the session, and participants were presented with the following questions: “Do you think the active stimulation was in the first session or the second session?”.

### Procedure

A double-blind, sham tDCS-controlled, repeated measurs design was used. Research sessions were carried out in custom brain stimulation laboratories at the University of Kent. Participants, received either the active (anodal) or placebo (sham) focal tDCS in two sessions with a minimum 72-hour interval between sessions at left IFG or left TPJ regions. Participants completed a larger language battery consisting of picture naming and verbal fluency tasks during both sessions while they were getting active or sham stimulation. We analysed and focused on the fluency task results within this study as the naming results have been published elsewhere (Yucel et al, 2025). In the first session, prior to the stimulation and language battery, all participants completed the matrix reasoning task (MaRs-IB) as a measure of fluid intelligence.

### Statistical Analyses

We conducted a 2×2×2×2×2 repeated-measures ANOVA to assess whether sham and anodal stimulation (STIMULATION TYPE) to the left IFG or left TPJ (REGION) altered semantic and phonemic fluency (FLUENCY TYPE) performance in the first 15 seconds versus the final 45 seconds (TIME) in healthy younger and older adults (AGE GROUP). We conducted a follow-up analysis adding in fluid intelligence as a covariate to assess whether general cognitive performance predicted stimulation response.

To evaluate the effectiveness of blinding, chi-squared tests were conducted to examine associations between participants’ ability to identify the active stimulation session and the factors of age group (younger vs. older) and stimulated region (left IFG vs. left TPJ).

For mood outcomes, Visual Analogue Mood Scale (VAMS) scores were analysed separately for positive and negative mood using a 2 (Stimulation: Anodal vs. Sham) × 2 (Age Group: Younger vs. Older) × 2 (Region: Left IFG vs. Left TPJ) repeated measures ANOVA, with stimulation as the within-subjects factor and age group and region as between-subjects factors. Partial eta squared (η²p) was reported as a measure of effect size.

Adverse effects were assessed using the same 2 × 2 × 2 repeated measures ANOVA structure to compare anodal and sham sessions across age groups and regions.

An alpha level of .05 (two-tailed) was used to determine statistical significance for all analyses.

## Results

Performance on both phonemic and semantic fluency across both sham and anodal stimulation sessions at the left IFG and left TPJ are presented in Table 2.

**Table 2.** Mean number of words produced in phonemic and semantic fluency tasks (summed across easy and hard conditions) under sham and anodal stimulation at the left inferior frontal gyrus (IFG) and left temporoparietal junction (TPJ).

| Young Adults | Left IFG |  | Left TPJ |  |
| --- | --- | --- | --- | --- |
|  | sham | anodal | sham | anodal |
| Phonemic |  |  |  |  |
| Fluency |  |  |  |  |
| *1-15 secs* | 9.69 (3.19) | 9.97 (3.17) | 9.67 (3.61) | 9.08 (3.00) |
| *15-60 secs* | 14.53 (4.75) | 14.03 (5.14) | 13.94 (5.72) | 15.89 (5.37) |
| Semantic |  |  |  |  |
| Fluency |  |  |  |  |
| *1-15 secs* | 16.97 (3.46) | 17.67 (4.22) | 16.19 (2.90) | 16.39 (2.82) |
| *15-60 secs* | 19.78 (7.06) | 19.75 (6.06) | 20.97 (5.64) | 20.17 (6.35) |
| Older Adults | Left IFG |  | Left TPJ |  |
|  | sham | anodal | sham | Anodal |
| Phonemic |  |  |  |  |
| Fluency |  |  |  |  |
| *1-15 secs* | 11.14 (3.05) | 11.17 (2.82) | 10.00 (2.92) | 10.86 (2.66) |
| *15-60 secs* | 19.42 (7.44) | 16.72 (6.70) | 17.33 (5.55) | 18.19 (4.39) |
| Semantic |  |  |  |  |
| Fluency |  |  |  |  |
| *1-15 secs* | 15.67 (3.23) | 16.47 (3.33) | 15.58 (3.24) | 15.56 (3.37) |
| *15-60 secs* | 21.06 (5.36) | 22.08 (6.73) | 20.19 (5.55) | 21.03 (7.03) |

### Main Effects and Higher-Order Interactions

A significant main effect of fluency type was observed, *F*(1,140) = 301.15, *p* < .001, η² = .683, indicating overall differences in performance between phonemic and semantic tasks. This effect was qualified by a significant Age Group × Fluency Type interaction, *F*(1,140) = 14.30, *p* < .001, η² = .093, suggesting differential effects of fluency type across age groups.

A main effect of time was also identified, *F*(1,140) = 335.81, *p* < .001, η² = .706, indicating that fluency performance varied over time. This effect was qualified by an Age Group × Time interaction, *F*(1,140) = 12.74, *p* < .001, η² = .083, and a Fluency Type × Time interaction, *F*(1,140) = 13.02, *p* < .001, η² = .085.

Importantly, a Stimulation Type × Fluency Type × Region interaction was identified, *F*(1,140) = 6.77, *p* = .010, η² = .046, as well as a Stimulation Type × Time × Region interaction, *F*(1,140) = 4.96, *p* = .028, η² = .034. A four-way interaction between Stimulation Type, Fluency Type, Time, and Age Group was also significant, *F*(1,140) = 5.63, *p* = .019, η² = .039. No other interactions involving stimulation type were significant (*ps* between .067 and .914).

### Stimulation Type × Fluency Type × Region

To investigate the three-way interaction involving fluency type and brain region, separate analyses were conducted for phonemic and semantic fluency.

In phonemic fluency, the Stimulation Type × Region interaction was significant, *F*(1,142) = 8.05, *p* = .005, η² = .054. In the inferior frontal gyrus (IFG), the main effect of stimulation type did not reach significance, *F*(1,71) = 3.61, *p* = .061, η² = .048. However, in the temporoparietal junction (TPJ), a significant effect of stimulation type was observed, *F*(1,71) = 4.49, *p* = .038, η² = .059, with higher fluency scores following anodal compared to sham stimulation.

In semantic fluency, the Stimulation Type × Region interaction was not significant, *F*(1,142) = 0.77, *p* = .381, η² = .005. These findings suggest that the regional effects of stimulation were specific to phonemic fluency. Figure X illustrates the effects of stimulation across regions.

**Figure X.**
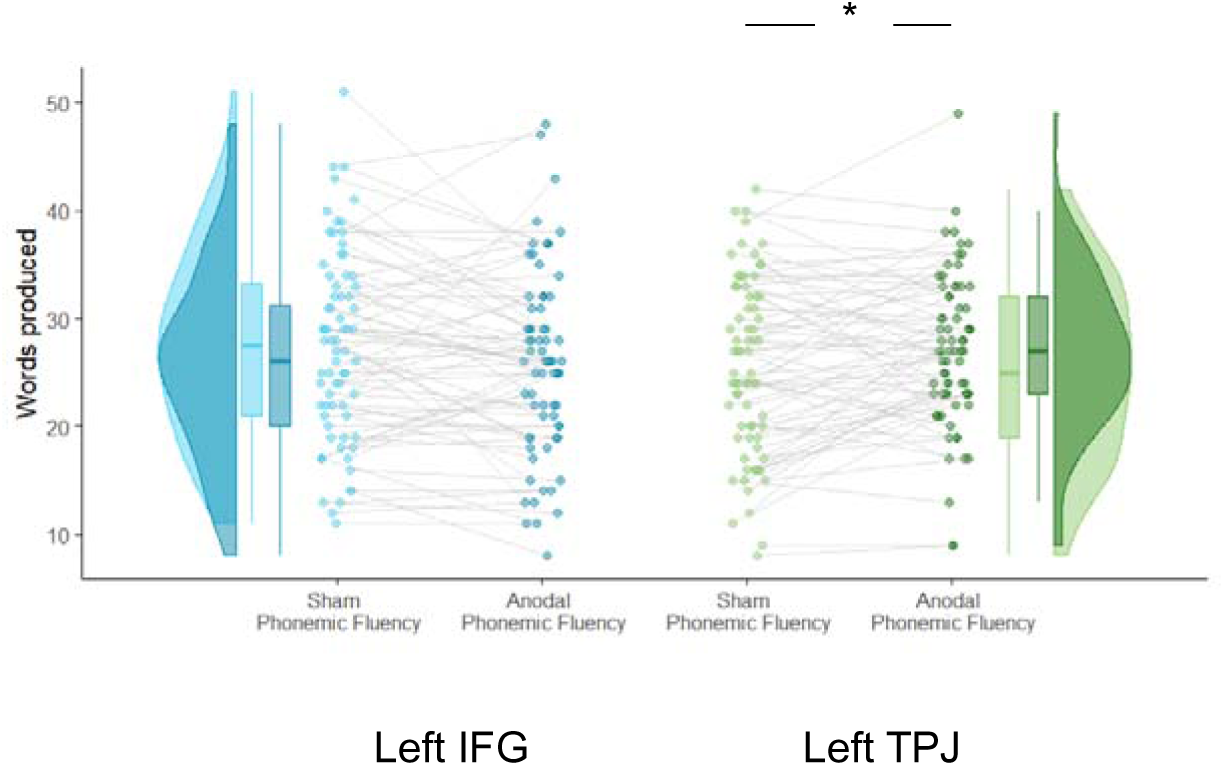
Stimulation effects on phonemic fluency at both the left IFG and TPJ. *\* p< .05*

### Stimulation Type × Time × Region

To explore the interaction between stimulation, time, and region, we conducted separate analyses for the IFG and TPJ.

In the IFG, the Stimulation Type × Time interaction was significant, *F*(1,71) = 4.00, *p* = .049, η² = .053. However, follow-up comparisons revealed that stimulation type did not significantly affect word generation in either the initial 15 seconds, *F*(1,71) = 2.50, *p* = .118, or the final 45 seconds, *F*(1,71) = 1.15, *p* = .288.

In the TPJ, the Stimulation Type × Time interaction was not significant, *F*(1,71) = 1.38, *p* = .245, η²L = .019. These results indicate that stimulation timing effects were more prominent in the IFG. Figure X depicts fluency performance over time across both regions.

**Figure X.**
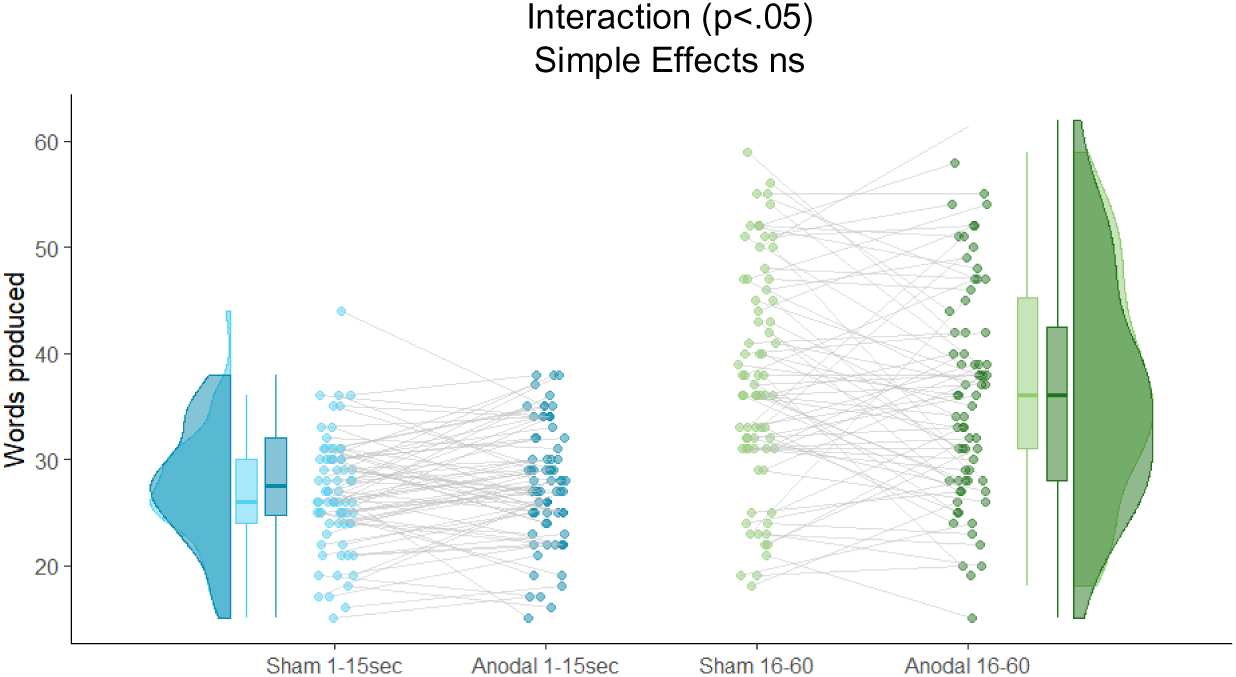
Stimulation effects across the first 15 seconds and last 45 seconds on combined phonemic and semantic fluency. *Note.* A significant interaction was identified showing that anodal-tDCS to the left IFG increased the number of words produced in the first 15 seconds but decreased the number of words produced in the final 45 seconds.

### Stimulation Type × Fluency Type × Time × Age Group

To examine the four-way interaction involving fluency type, time, and age group, we analysed semantic and phonemic fluency tasks separately.

In semantic fluency, the Stimulation Type × Time × Age Group interaction was not significant, *F*(1,142) = 2.52, *p* = .115, η²L = .017. In phonemic fluency, however, the interaction was significant, *F*(1,142) = 5.61, *p* = .019, η² = .038.

Follow-up analyses within each age group indicated that the Stimulation Type × Time interaction was not significant in either older adults, *F*(1,71) = 3.50, *p* = .066, η² = 0.05, or younger adults, *F*(1,71) = 2.11, *p* = .150, η² = 0.03.

The Age Group x stimulation type interaction was not significant in the first 15 seconds, *F*(1,142) = 1.48, *p* = .226, η² = 0.01, or the final 45 seconds, *F*(1,142) = 3.42, *p* = .067, η² = 0.02.

The Age Group x Time interaction was significant during sham stimulation, F(1,142)=15.18, p<.001, η² = 0.10 but not during anodal stimulation, F(1,142)=1.84, p=.18, η² = 0.01.

These findings suggest that the overall four-way interaction was driven by phonemic fluency, where the effects of stimulation varied subtly over time and differed by age group. Specifically, anodal stimulation over the left IFG selectively disrupted older adults’ ability to sustain phonemic fluency over time, reducing their later word generation relative to sham.

### Moderation by Fluid Intelligence

To determine whether fluid intelligence moderated stimulation responses, we conducted a repeated-measures ANOVA with MaRs-IB scores as a covariate. A significant Stimulation Type × Region × MaRs-IB interaction was found, *F*(1,138) = 5.40, *p* = .022, η² = .038, indicating that individual differences in fluid intelligence influenced regional effects of stimulation.

No other interactions involving MaRs-IB were significant. Specifically, the Stimulation Type × MaRs-IB interaction was not significant, *F*(1,138) = 1.26, *p* = .265, η² = .009, nor was the Stimulation Type × Time × MaRs-IB interaction, *F*(1,138) = 0.97, *p* = .326, η² = .007.

Follow-up analyses by region showed that in the TPJ, a significant Stimulation Type × MaRs-IB interaction emerged, *F*(1,70) = 7.80, *p* = .007, η² = .100, whereas no such interaction was found in the IFG, *F*(1,70) = 0.82, *p* = .368, η² = .012.

**Figure X.**
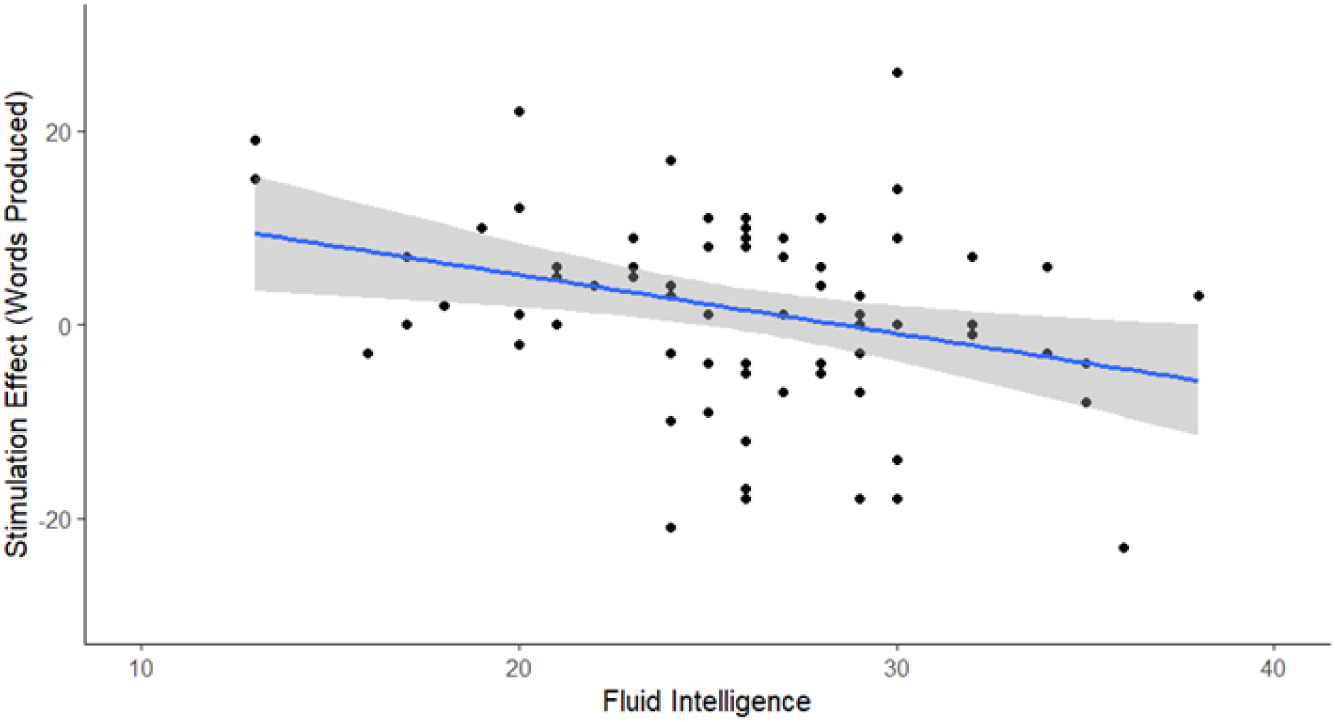
Scatterplot displaying the relationship between fluid intelligence and stimulation effect on phonemic and semantic fluency combined at the left TPJ

### Blinding Effect

The effectiveness of blinding was demonstrated by the use of a chi-squared test to examine the impact of participants’ age group and stimulated region on their ability to identify the active session accurately. The chi-squared test showed a statistically significant association between the age group and blinding scores (χ^2 = 6.399, df = 1, p = 0.011), indicating that age group affected the participants’ ability to maintain blinding integrity consistently. Compared to older individuals, younger adults had a superior ability in predicting active sessions. However, no significant association was discovered between the stimulated area and blinding scores (χ^2 = 0.256, degrees of freedom = 1, p > 0.05).

### Positive Mood

A 2x2x2 repeated measures ANOVA was conducted to assess the impact of Anodal and Sham sessions on VAMS positive mood scores in older and younger individuals for left IFG and left TPJ. A stimulation effect was observed, with statistical significance (F(1, 140) = 4.934, p = .028, η²p = 0.034). Participants reported a more substantial reduction in VAMS positive scores following anodal stimulation (M = -10.82) compared to the sham stimulation (M = - 2.80). The study found no significant interaction between STIMULATION X AGE GROUP (F(1,140) = 0.042, p = .838, η²p = .000) and STIMULATION X REGION (F(1,140) = 0.035, *p* = .852, η²p = .000) nor between STIMULATION X AGE GROUP X REGION (F(1,140) = 3.224, p = .075, η²p = .023).

### Negative Mood

The study found no significant stimulation effect on VAMS negative scores, *F*(1, 140) = 0.702, p = .404, η²p = 0.005. The impact of stimulation on VAMS scores was found to be nonsignificant when considering the other interactions of STIMULATION X AGE GROUP *F*(1,140) < 0.001, *p* = .993, η²p < .001, STIMULATION X REGION *F*(1,140) = 0.068, *p* = .795, η²p < .001, and STIMULATION X AGE GROUP X REGION *F*(1,140) = 0.171, *p* = .680, η²p = .001.

### Adverse Effects

The participants reported significantly higher adverse impact scores for the anodal session (*M* = 1.88) compared to the sham session (*M* = 1.58). However, the stimulation effect was not statistically significant, *F*(1,140) = 3.451, *p* = .065, η*²p* = .024. There were no significant interaction effects seen between the stimulation type and age group *F*(1,140) = 0.423, *p* = .517, η*²p* = .003, the stimulation type and region *F*(1,140) = 0.009, *p* = .926, η*²p* < .001, and the stimulation type and age group and region, *F*(1,140) = 0.863, *p* = .355, η*²p* = .006.

## Discussion

We investigated the effects of focal anodal transcranial direct current stimulation (tDCS) on phonemic and semantic verbal fluency in younger and older adults, targeting the left inferior frontal gyrus (IFG) and the left temporoparietal junction (TPJ). We examined how stimulation site, fluency type, energization, and individual differences in fluid intelligence shaped stimulation effects.

A key finding was that stimulation response varied with fluid intelligence. Greater gains in total fluency scores following left TPJ stimulation were observed in individuals with lower baseline fluid intelligence, whereas higher ability was associated with smaller or even detrimental effects (Perceval et al., 2020). This baseline-dependent pattern supports the state-dependency framework, which holds that neuromodulation efficacy depends on the initial functional state of the targeted network (Silvanto et al., 2008). tDCS alters cortical excitability by shifting resting membrane potentials and enhancing synaptic efficacy, particularly in underactive or suboptimally engaged circuits (Stagg & Nitsche, 2011). For individuals with lower fluid intelligence, language-related networks such as the TPJ and its connections to frontal cortex may operate below optimal efficiency (Jung & Haier, 2007), leaving greater capacity for stimulation to enhance plasticity and task-related activation (Meinzer et al., 2013). By contrast, high-functioning individuals may already operate close to their optimal range, leaving little room for improvement and increasing the risk of overstimulation (Santarnecchi et al., 2015). These findings align with evidence that baseline cognitive performance shapes responsiveness to stimulation (Yucel et al., 2025; Benwell et al., 2015; Berryhill & Jones, 2012).

We also found that stimulation of the left TPJ significantly enhanced phonemic fluency, whereas stimulation of the left IFG did not. This is noteworthy, as posterior temporal and temporoparietal regions are typically linked with semantic processing and comprehension, while phonemic fluency is more closely tied to frontal executive control (Birn et al., 2010; Hirshorn & Thompson-Schill, 2006). Yet phonemic fluency also depends on access to phonological representations and the temporary storage and manipulation of verbal material, processes supported by the dorsal language stream and the TPJ (Hickok & Poeppel, 2007; Biesbroek et al., 2021). Stimulation here may therefore enhance retrieval by strengthening phonological processing and working memory (Buchsbaum et al., 2001; Indefrey & Levelt, 2004). This suggests that posterior language regions, although less traditionally associated with fluency, can play a critical role in supporting phoneme-based word generation.

At the IFG, stimulation influenced the *timing* rather than the total amount of fluency output. Participants generated slightly more words in the first 15 seconds under anodal stimulation, but fewer in the subsequent 45 seconds. This pattern indicates a transient benefit for verbal response initiation, followed by reduced sustained output. The left IFG has been linked to energization—the ability to initiate and maintain effortful activity (Cipolotti et al., 2021; Stuss & Alexander, 2007)—and stimulation may have briefly boosted these initiation-related processes, consistent with evidence that this region supports goal activation and response coordination under time pressure (Sharp et al., 2010; Ghanavati et al., 2019). Although modest, these results suggest a link between IFG stimulation, energization, and early fluency output, but at the cost of sustained responding.

Importantly, older adults showed a decline in phonemic fluency during the final 45 seconds of anodal stimulation relative to sham, suggesting that stimulation interfered with processes that normally help maintain word retrieval under increasing task demands. This effect was most evident following stimulation of the left IFG. Although anodal tDCS is often assumed to facilitate performance (Jacobson et al., 2012), the present results may reflect disruption of compensatory recruitment mechanisms in prefrontal cortex that older adults rely on during effortful word generation (Cabeza et al., 2002; Reuter-Lorenz & Cappell, 2008). By contrast, semantic fluency was unaffected, consistent with its lesser reliance on executive control (Henry & Crawford, 2004; Shao et al., 2014).

Secondary findings provide further context. Younger adults were more accurate in identifying stimulation conditions, echoing age-related differences in sensory perception (Gandiga et al., 2006). Stimulation also slightly reduced positive affect, in line with prior reports of mood modulation by tDCS (Brunoni et al., 2014). Adverse effects were mild and consistent with established safety guidelines (Antal et al., 2017).

This study has several strengths, including preregistration and a targeted stimulation design, but limitations should be acknowledged. Effects were generally modest with notable inter-individual variability, consistent with findings in older adults (Fromm et al., 2025; Vergallito et al., 2022). Our single-session design restricts conclusions about longer-term effects; future research should test multisession interventions and examine dose–response relationships, ideally integrating neurocognitive and neuroimaging profiles to predict responsiveness (Filmer et al., 2020; Fromm et al., 2025; Niemann et al., 2024). Older adults also outperformed younger adults on fluency tasks, likely due to higher educational attainment, limiting generalisability. Greater sampling diversity will be important in future work.

In conclusion, this study demonstrates that focal tDCS can modulate verbal fluency performance depending on stimulation site, task demands, and baseline cognitive ability. The findings highlight that stimulation effects on language are not uniform: they reflect the interaction between individual cognitive state, temporal dynamics of task performance, and the specific processes engaged at different cortical sites. Future work using multisession and individualised stimulation protocols will be essential to clarify the potential of tDCS for enhancing fluency in both healthy and pathological ageing.

## Data Availability

Data will be available online at https://osf.io/67g859

